# S-acylation stabilizes ligand-induced receptor kinase complex formation during plant pattern-triggered immune signalling

**DOI:** 10.1101/2021.08.30.457756

**Authors:** Charlotte H. Hurst, Dionne Turnbull, Kaltra Xhelilaj, Sally Myles, Robin L. Pflughaupt, Michaela Kopischke, Paul Davies, Susan Jones, Silke Robatzek, Cyril Zipfel, Julien Gronnier, Piers A. Hemsley

## Abstract

Plant receptor kinases are key transducers of extracellular stimuli, such as the presence of beneficial or pathogenic microbes or secreted signalling molecules. Receptor kinases are regulated by numerous post-translational modifications. Here, using the immune receptor kinases FLS2 and EFR, we show that S-acylation at a cysteine conserved in all plant receptor kinases is crucial for function. S-acylation involves the addition of long-chain fatty acids to cysteine residues within proteins, altering their biophysical properties and behaviour within the membrane environment. We observe S-acylation of FLS2 at C-terminal kinase domain cysteine residues within minutes following perception of its ligand flg22, in a BAK1 co-receptor dependent manner. We demonstrate that S-acylation is essential for FLS2-mediated immune signalling and resistance to bacterial infection. Similarly, mutating the corresponding conserved cysteine residue in EFR supressed elf18 triggered signalling. Analysis of unstimulated and activated FLS2-containing complexes using microscopy, detergents and native membrane DIBMA nanodiscs indicates that S-acylation stabilises and promotes retention of activated receptor kinase complexes at the plasma membrane to increase signalling efficiency.

## Introduction

The plasma membrane defines the boundary between the cell interior and the external environment. Receptor kinases (RKs) found in the plasma membrane act as the principle means of perception for most of the stimuli that a plant encounters, such as hormones, signalling peptides and microbe associated molecular patterns (MAMPs). RKs comprise the largest single gene family in plants [1, 2] and are central to current efforts to breed or engineer crops able to withstand emerging pathogen threats, interact with beneficial microbes or better tolerate abiotic stress [3–6]. Understanding the mechanisms and principles underlying the formation and activation of RK complexes is therefore critical to informing these approaches.

The RK FLAGELLIN SENSING 2 (FLS2) is the receptor for bacterial flagellin and the flagellin-derived peptide flg22 [7], and is an archetype for RK research, particularly in the area of host-microbe interactions. flg22 binding to the extracellular leucine-rich repeats of FLS2 induces interaction with the co-receptor BAK1/SERK3 (BRI1-ASSOCIATED RECEPTOR KINASE 1/SOMATIC EMBRYOGENESIS RECEPTOR-LIKE KINASE 3). Subsequent transphosphorylation of FLS2 by BAK1 initiates a cascade of immune signalling to activate anti-bacterial defence responses. As part of this overall process, flg22 perception leads to changes in FLS2 phosphorylation [8], SUMOylation and ubiquitination [10] state, indicating a high degree of post-translational regulation. FLS2 activation also alters overall complex composition [7, 9–18] and physical properties [19]. However, the underlying mechanisms and functional relevance of these changes remain unknown.

S-acylation is a reversible post-translational modification, whereby long chain fatty acids are added to cysteine residues by protein S-acyl transferases [20] and removed by acyl-protein thioesterases [21]. This modification can lead to changes in protein trafficking, stability, and turnover. S-acylation has been proposed to drive membrane phase partitioning [22, 23] while changes in protein S-acylation state have been hypothesised to modulate protein-protein and protein-membrane interactions, or even alter protein activation states [24], direct experimental evidence to support these ideas is lacking. We recently discovered that FLS2, alongside all other plant RKs tested, is post-translationally modified by S-acylation [25]. Here we demonstrate that ligand induced S-acylation of FLS2 and EFR RKs, at a site conserved in all RKs across the span of plant evolutionary history, acts as a positive regulator of signal transduction. Mechanistically, ligand-induced S-acylation of FLS2 promotes stability and retention of receptor complexes at the plasma membrane.

## Results

### FLS2 undergoes ligand responsive S-acylation

Our previous analysis of FLS2 identified the juxta-transmembrane (TM) domain cysteines (Cys830,831) as being constitutively S-acylated, but this modification was dispensable for FLS2 function [25]. All RK superfamily members subsequently tested, with or without a juxta-TM S-acylation site homologous to FLS2 C^830,831^, also appear to be S-acylated [25]. This indicates that non-juxta-TM S-acylation sites, potentially conserved in all RKs, exist. Other post-translational modifications affecting FLS2, and the broader RK superfamily, including phosphorylation [26], ubiquitination [10] and SUMOylation [9] are all responsive to ligand binding. Given the dynamic nature of S-acylation [21] we were interested to determine whether FLS2 S-acylation state is also ligand responsive. In Col-0 wild type plants, FLS2 S-acylation significantly increased above basal levels in controls following 20-min exposure to the FLS2 agonist peptide flg22. FLS2 S-acylation subsequently returned to basal levels within 1 h (figures 1A and 1B). Consistent with its ligand-dependency, FLS2 S-acylation was contingent upon the FLS2 co-receptor BAK1 (BRI1-ASSOCIATED KINASE) (figure 1C). PUB12/13 (PLANT U-BOX12/13) are ubiquitin ligases proposed to promote FLS2 endocytosis and attenuate signalling [10]. FLS2 S-acylation is impaired in the *pub12/13* double mutant, suggesting that PUB12/13 action may be required for S-acylation to occur. Additionally, flg22 induced S-acylation of FLS2 was unaffected in *chc2-1* (CLATHRIN HEAVY CHAIN 2) mutants [17] of clathrin heavy chain 2 (figure 1C). These data indicate that FLS2 S-acylation occurs after initiation of FLS2 signalling and the hypothesised ubiquitination thought to mark FLS2 for internalisation, but before endocytosis of FLS2 occurs. Treatment of Arabidopsis Col-0 plants with elf18, an immunogenic peptide derived from bacterial elongation factor Tu, recognised by the RK EFR (ELONGATION FACTOR-Tu RECEPTOR) that acts similarly to FLS2 [27], failed to elevate FLS2 S-acylation (figure 1D). This demonstrates that the increase in FLS2 S-acylation is specifically linked to activation of FLS2 signalling and not a general phenomenon related to activation of RK-mediated defence responses.

**Figure 1.**
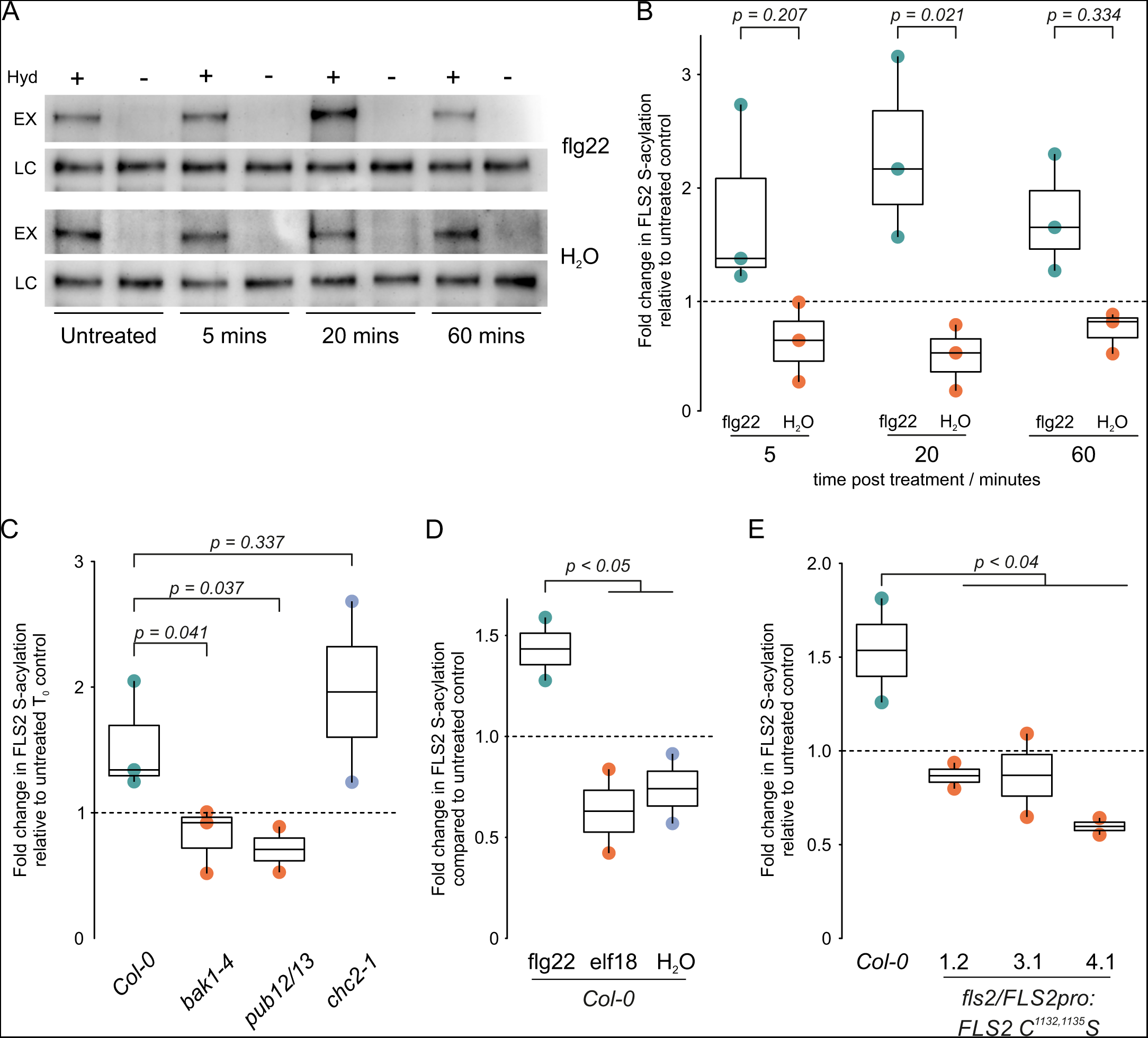
FLS2 S-acylation increases upon flg22 perception. **A**. Representative western blot of FLS2 S-acylation state in Arabidopsis Col-0 plants treated with 1 μM flg22 peptide or water as determined by acyl-biotin exchange assay. EX - indicates S-acylation state, LC - loading control, Hyd - indicates presence (+) or absence (−) of hydroxylamine. **B**. Quantification of western blot data in A. showing change in S-acylation state in Arabidopsis *Col-0* plants treated with 1 μM flg22 (green) or water (orange). S-acylation state is shown relative to untreated plants (black dashed line). n = 3 biological repeats. Box plot shows median and IQR, whiskers indicate data points within 1.5 × IQR. Significance of difference between flg22 and water treatments at each timepoint was determined by ANOVA and Tukey’s HSD test. **C**. S-acylation of FLS2 in response to flg22 requires BAK1 and PUB12/13 but not CHC2. S-acylation state was determined by acyl-biotin exchange after 20 minutes exposure to 1 μM flg22 and is shown relative to untreated Arabidopsis plants of the same genotype (dashed line). Box plot shows median and IQR, whiskers indicate data points within 1.5 × IQR. Significant differences of each genotype to flg22 treated Arabidopsis *Col-0* as determined by Student’s t-test are shown. **D**. FLS2 undergoes S-acylation in response to flg22 treatment but not elf18. S-acylation state as determined by acyl-biotin exchange after 20 minutes of treatment using 1 μM peptide or water is shown relative to untreated Arabidopsis plants (black, dashed line). Box plot shows median and IQR, whiskers indicate data points within 1.5 × IQR. Significant differences of elf18 or water treatment compared to flg22 treated Arabidopsis *Col-0* as determined by Student’s t-test are shown. **E**. FLS2 C^1132,1135^S mutants are blocked in flg22 mediated increases in S-acylation. S-acylation state is shown following 20 minutes 1 μM flg22 treatment relative to untreated Arabidopsis plants of the same genotype (black, dashed line). Box plot shows median and IQR, whiskers indicate data points within 1.5 × IQR. Significant difference of each line compared to flg22 treated *Col-0* as determined by Student’s t-test are shown.

### flg22 responsive S-acylation sites of FLS2 are located in the kinase domain C-terminus and are conserved across the wider plant receptor kinase superfamily

FLS2 C^830,831^S mutants [25] lacking the juxta-TM S-acylation retain the ability to be S-acylated in response to flg22 (figure S1A, 5 and 20 minute timepoints). FLS2 therefore contains S-acylation sites in addition to C^830,831^ that are responsive to ligand perception. While FLS2 C^830,831^S expressed at native levels in unstimulated Arabidopsis is very weakly S-acylated [25] (figure S1A, untreated), we observed S-acylation of FLS2 C^830,831^S in the absence of flg22 when overexpressed in *Nicotiana benthamiana*. Mutation of FLS2 Cys 1132 and 1135 in addition to Cys 830 and 831 (FLS2 C^830,831,1132,1135^S) abolished FLS2 S-acylation compared to FLS2 C^830,831^S (figure S1B) when overexpressed in *N. benthamiana*, suggesting that Cys1132/1135 are sites of S-acylation. Following this observation, we found that *fls2c/proFLS2:FLS2 C^1132,1135^S* Arabidopsis plants (figure S1C) showed no increase in S-acylation following flg22 treatment (figure 1E), indicating that these cysteines are the sites of ligand inducible S-acylation. Interestingly, 1-2 conserved cysteine residues at the C-terminus of the kinase domain corresponding to FLS2 Cys 1132 and/or 1135, are found across all RKs in Arabidopsis and RKs from Charophycean algae (figure S2) at the base of the broader Streptophyte lineage with high quality genome assemblies [28], suggesting a evolutionarily conserved and important role for these cysteines. In support of this hypothesis, we found that EFR-GFP [29] transiently expressed in *N. benthamiana* undergoes an elf18-induced increase in S-acylation, and this is blocked by mutation of the EFR Cys 975, the cysteine homologous to flg22 responsive FLS2 Cys 1135 (figure S1D, S1E, S2).

### Receptor kinase C-terminal S-acylation is required for early immune signalling

Consistent with the evolutionarily conserved nature of the FLS2 kinase domain S-acylated cysteines amongst RKs, mutation of these cysteines affects FLS2 function. *fls2c/proFLS2:FLS2 C^1132,1135^S* plants are impaired in several aspects of early immune signalling, such as reactive oxygen species production, MAP kinase activation and immune gene expression (figure 2A, B, C). In the absence of flg22 ligand, both FLS2-3xMyc-GFP and FLS2 C^1132,1135^S-3xMyc-GFP show similar accumulation at the plasma membrane (figure S3A; water treatments) and similar lateral membrane mobility (figure S3B, C; water treatments). Remorins are plasma membrane resident proteins that form clusters and have been proposed as markers for membrane nanodomains [30]. Specifically, REM1.3 (Remorin 1.3) nanodomains have previously been shown to have strong spatial overlap with FLS2 nanodomains [31]. Both FLS2-3xMyc-GFP and FLS2 C^1132,1135^S-3xMyc-GFP show similar co-localization with REM1.3 nanodomains (figure S3D, E, F). These data combined indicate that there is no aberrant basal cellular behaviour of the FLS2 C^1132,1135^S mutant when compared to FLS2 that could account for the observed reduction in response to flg22. To determine whether the conserved C-terminal cysteines may have a general role in RK function, we transiently expressed EFR-GFP [29] and EFR C^975^S-GFP in *N. benthamiana*. We observed that elf18-induced MAP kinase phosphorylation and immune gene induction was reduced in EFR C^975^S-GFP expressing plants compared to EFR-GFP (figure 2D, E). This demonstrates that mutation of the conserved C-terminal cysteine in both FLS2 and EFR has a similar effect on early outputs and indicates a conserved mode of action. Structural homology modelling of FLS2 indicates that the C^1132,1135^S mutation does not affect FLS2 kinase domain structure (figure S4). Kinase activity is also dispensable for activation of signalling by EFR [32]. The observed effects of the FLS2 C^1132,1135^S and EFR C^975^S mutations on early signalling therefore cannot be readily explained through deleterious effects on kinase activity or structure.

**Figure 2.**
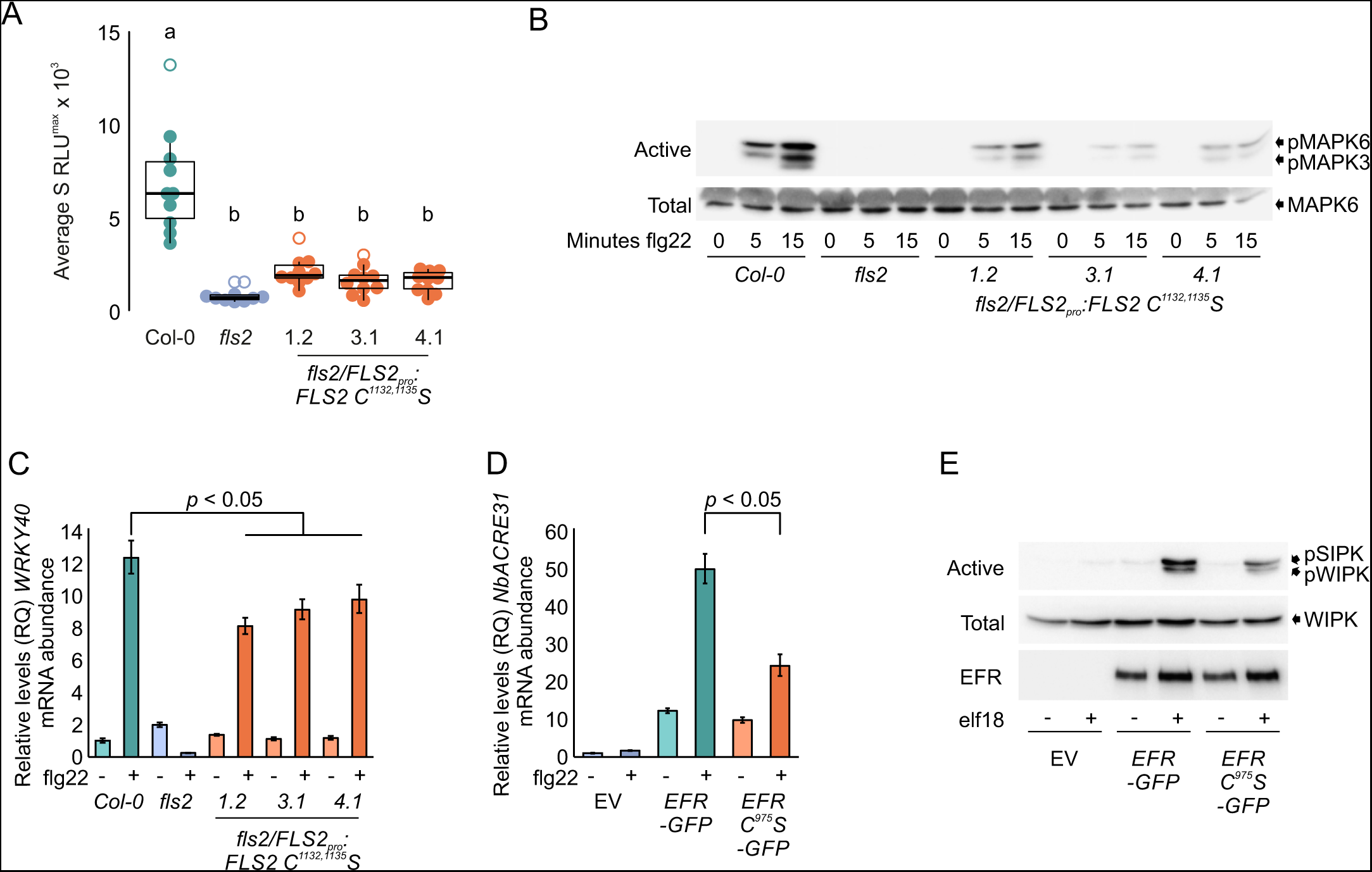
Acute responses to bacterial elicitor perception are reduced in FLS2 C^1132,1135^S and EFR-C^975^S expressing plants. **A**. ROS production induced by 100 nM flg22 treatment of Arabidopsis seedlings. Data points are the sum of the 3 highest consecutive readings per sample. n = 10 per genotype. Statistical outliers are shown as open circles. Box shows median and IQR, whiskers show +/− 1.5 × IQR. Statistically significant differences at p < 0.01 are indicated (a, b) and were calculated using ANOVA and Tukey HSD tests. **B**. MAPK activation in *fls2/FLS2pro:FLS2 C^1132,1135^S* Arabidopsis seedlings in response to 100 nM flg22 as determined over time by immunoblot analysis. pMAPK6/pMAPK3 show levels of active form of each MAPK. MAPK6 indicates total levels of MAPK6 as a loading control. Upper shadow band in MAPK6 blot is RuBisCO detected non-specifically by secondary antibody. **C**. *WRKY40* mRNA abundance after 1 hour treatment with 1 μM *flg22 in fls2/FLS2pro:FLS2 C^1132,1135^S* Arabidopsis seedlings as determined by qRT-PCR. Values were calculated using the ΔΔCT method, error bars represent RQMIN and RQMAX and constitute the acceptable error level for a 95% confidence interval according to Student’s t-test. **D**. *NbACRE31* mRNA abundance after 3 hour treatment with 1 μM elf18 in *EFR-GFP* and *EFR C^975^S-GFP* expressing *N. benthamiana* plants as determined by qRT-PCR. Values were calculated using the ΔΔCT method, error bars represent RQMIN and RQMAX and constitute the acceptable error level for a 95% confidence interval according to Student’s t-test. **E**. MAPK activation in *EFR-GFP* and *EFR C^975^S*-GFP expressing *N. benthamiana* plants in response to 15 minutes treatment with 1 μM elf18 as determined by immunoblot analysis. pSIPK/pWIPK show levels of active form of each MAPK. WIPK indicates total levels of WIPK as a loading control. EFR-GFP and EFR C^975^S-GFP levels are shown as a control for dosage effects on MAPK activation.

### FLS2 kinase domain S-acylation is required for late immune responses and anti-bacterial immunity

Early signalling outputs resulting from bacterial perception by FLS2 lead to longer term sustained responses to promote immunity. In line with decreased early immune responses, later flg22-induced gene expression and physiological outputs, such as *PR1* (PATHOGENESIS-RELATED GENE 1) expression and seedling growth inhibition, were affected in *fls2c/proFLS2:FLS2 C^1132,1135^S* plants (figure 3A, B). As a result of these cumulative signalling defects, FLS2 C^1132,1135^S failed to complement the hyper-susceptibility of *fls2* mutant plants to the pathogenic bacterium *Pseudomonas syringae* pv. tomato (*Pto*) DC3000 (figure 3C).

**Figure 3.**
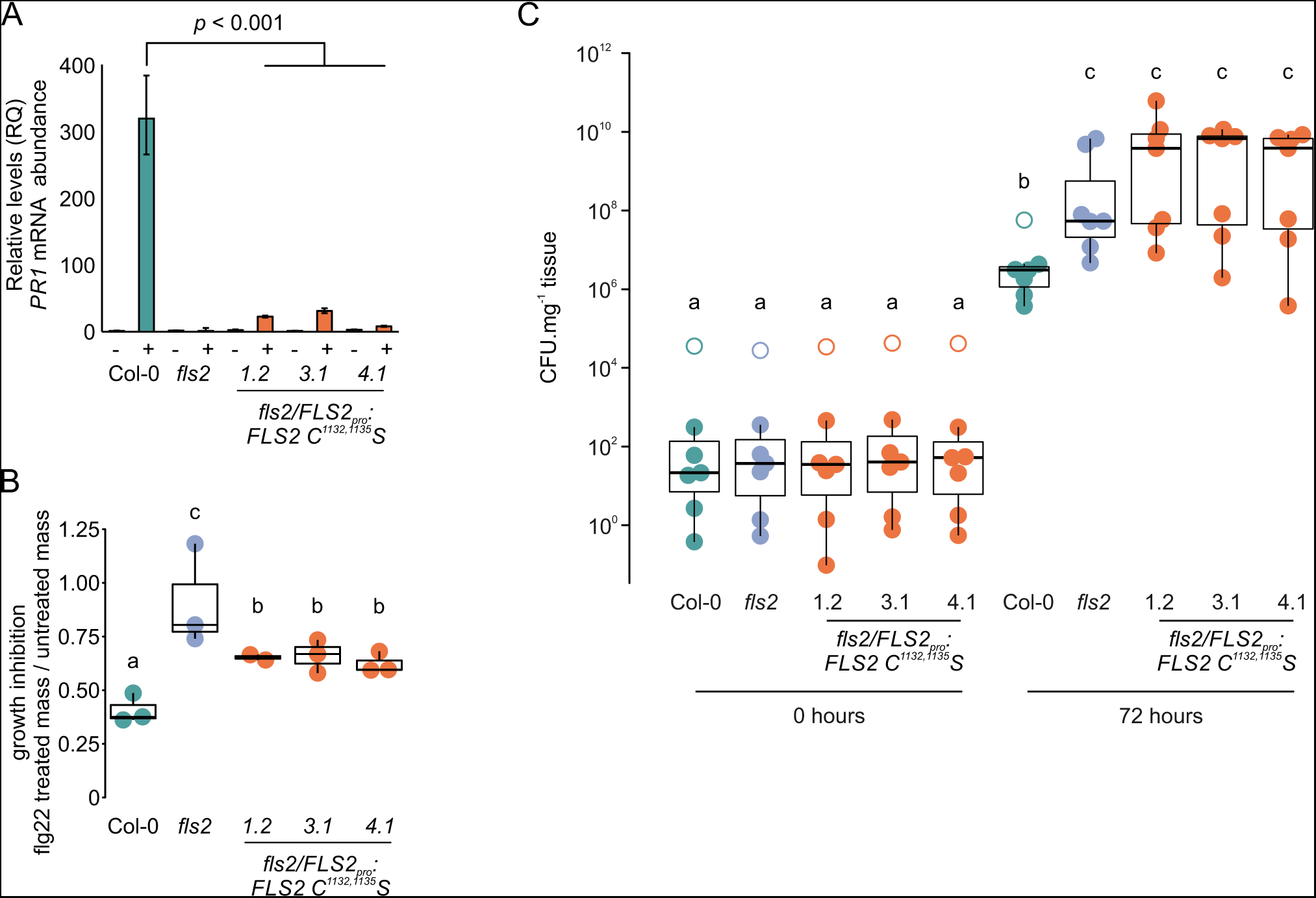
FLS2 S-acylation is required for long term immune response outputs. **A**. Induction of *PR1* gene expression after 24 hours treatment with 1 mM flg22 in *fls2/FLS2pro:FLS2 C^1132,1135^S* seedlings as determined by qRT-PCR. Values were calculated using the ΔΔ_CT_ method, error bars represent RQMIN and RQMAX and constitute the acceptable error level for a 95% confidence interval according to Student’s t-test. Significant differences in transcript mRNA detected in *fls2/FLS2pro:FLS2 C^1132,1135^S* Arabidopsis seedlings compared to Col-0 levels in flg22 treated samples are indicated. Similar data were obtained over 3 biological repeats. **B**. Inhibition of growth after 10 days of 1 μM flg22 treatment is reduced in fls2/FLS2pro:FLS2 C^1132,1135^S Arabidopsis seedlings. Box and whisker plots show data from 7 biological repeats (box denotes median and IQR, whiskers show +/− 1.5 × IQR), significant differences at p < 0.01 are indicated (a, b, c) and calculated by ANOVA with Tukey HSD test. **C**. Resistance to *P. syringae* pv. tomato DC3000 infection is impaired by loss of FLS2 S-acylation in *fls2/FLS2pro:FLS2 C^1132,1135^S* Arabidopsis plants. Box and whisker plots show data from 7 biological repeats (box denotes median and IQR, whiskers show +/− 1.5 × IQR, outliers are shown as open circles), significant differences at p < 0.05 are indicated (a, b, c) and calculated by ANOVA with Tukey HSD test.

### S-acylation of FLS2 stabilizes flg22-induced FLS2-BAK1 signalling complexes within the plasma membrane

Differential solubility in cold non-ionic detergents such as Triton X-100 or IGEPAL CA-630, leading to formation of detergent soluble or resistant membrane fractions (DSM and DRM respectively), has been used to characterise overall changes to protein physical properties, particularly in the context of protein S-acylation [19, 33]. *fls2c/proFLS2:FLS2* and *fls2c/proFLS2:FLS2 C^1132,1135^S* plants were treated with or without flg22 and total cold IGEPAL CA-630 protein extracts were separated into DRM and DSM/cytosol fractions [34]. Following flg22 treatment, FLS2 abundance in cold IGEPAL CA-630 derived DRMs showed a slight reduction, while FLS2 C^1132,1135^S DRM abundance decreased by ~50% (figure 4A, B). Overall, these data suggest a change in protein and/or lipid environment of the FLS2 C^1132,1135^S containing complex compared to wild type within 20 minutes of flg22 exposure.

**Figure 4.**
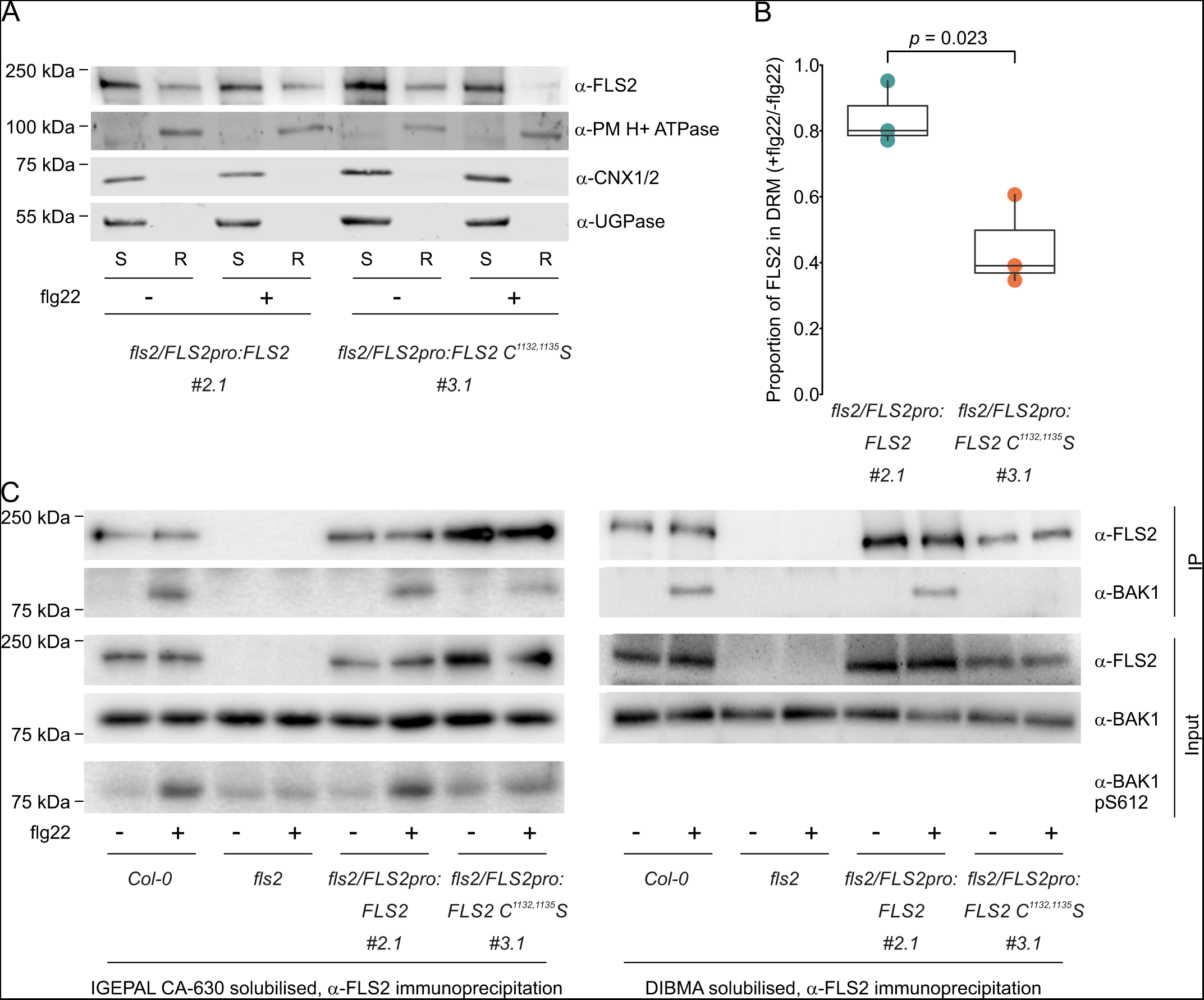
FLS2 C^1132,1135^S shows reduced interaction with BAK1 following flg22 stimulation. **A**. FLS2 C^1132,1135^S shown altered DRM partitioning compared to FLS2. Arabidopsis flg22 treated seedlings were lysed in cold IGEPAL CA-630 buffer and separated into detergent soluble (S) and detergent resistant (R) fractions. Relative partitioning of FLS2 into each fraction was determined by western blotting with anti-FLS2 rabbit polyclonal antibody. Purity of fractions is shown by western blot using anti-PM H+ ATPase (PM ATPase, DRM marker), anti-Calnexin1/2 (CNX1/2, DSM marker) and anti-UDP-glucose pyrophosphorylase (UGPase, cytosol marker) antibodies. **B**. Quantification of FLS2 data shown in A from 3 biological repeats. Box plot shows median and IQR, whiskers indicate data points within 1.5 × IQR. Significance was calculated using Student’s t-test. **C**. FLS2 was immunoprecipitated from IGEPAL CA-630 (left) or DIBMA (right) solubilised flg22 treated Arabidopsis seedling lysates using anti-FLS2 rabbit polyclonal antibody. BAK1 recovery was assessed using rabbit polyclonal anti-BAK1 antibody. flg22 induced BAK1 autophosphorylation at Ser612 was assessed in IGEPAL CA-630 solubilised input samples using rabbit polyclonal anti-BAK1 pS612 antibody.

Assessment of flg22-induced FLS2-BAK1 complex formation by co-immunoprecipitation following solubilisation with cold IGEPAL CA-630 [35] indicated that the observed FLS2-BAK1 interaction was reduced in FLS2 C^1132,1135^S mutants (figure 4C). Furthermore, flg22-induced BAK1 S^612^ auto-phosphorylation [36], used as a marker of *in vivo* complex formation, was also weaker in FLS2 C^1132,1135^S-expressing plants (figure 4C), supporting these biochemical observations. In contrast to IGEPAL CA-630, diisobutylene/maleic acid (DIBMA) copolymer does not form DRM-like fractions. DIBMA disrupts lipid-lipid, but not protein-protein or protein-lipid, interactions to form membrane nanodiscs containing protein complexes within their membrane environment [37]. Co-immunoprecipitation of DIBMA-solubilised FLS2-BAK1 and FLS2 C^1132,1135^S-BAK1 complexes after 20 minutes of flg22 treatment (figure 4C) indicates that FLS2-BAK1 interactions are likely stabilized by protein-protein and protein-lipid interactions that are reduced or absent from FLS2 C^1132,1135^S-BAK1 complexes.

Examination of FLS2 mobility by VA-TIRF microscopy (figure S3B, C) shows no detectable change in FLS2-3xMyc-GFP or FLS2 C^1132,1135^S-3xMyc-GFP motion within the plasma membrane following flg22 treatment. However, we observed a decrease in the number of particles of FLS2 C^1132,1135^S-3xMyc-GFP, but not wild type FLS2-3xMyc-GFP, at the plasma membrane following 20 minutes of flg22 treatment (figure S3B, C), suggesting premature or accelerated endocytosis of flg22 bound FLS2 C^1132,1135^S-3xMyc-GFP. Altogether, our observations indicate that FLS2 S-acylation stabilises FLS2-BAK1 association and maintains FLS2 in an signalling competent state at the plasma membrane.

## Discussion

FLS2, a prototypical RK, has been previously shown to be S-acylated at a pair of juxta-transmembrane domain cysteines (Cys 830,831), but S-acylation at these sites is apparently dispensable for function [25]. Here we demonstrate that FLS2 is S-acylated at additional cysteine residues (Cys 1132,1135) in a ligand-responsive manner and that this is required for efficient flg22-triggered signalling and resistance to *P. syringae* DC3000 bacterial infection. FLS2 S-acylation occurs within minutes of flg22 perception and requires the co-receptor BAK1 and the PUB12/13 ubiquitin ligases, but does not require CHC2 function (figure 1). We therefore propose that FLS2 S-acylation occurs as a result of FLS2 activation but precedes entry into the endocytic pathway. Supporting this hypothesis, preventing ligand mediated FLS2 S-acylation from occurring by using *fls2c/proFLS2:FLS2 C^1132,1135^S* plants reduces early signalling outputs, such as the phosphorylation of MAPK and the production of ROS (figure 2), processes unimpaired in mutants affecting FLS2 endocytosis [15, 17]. Indeed, our data (figure S3B, S3C) suggests a model where FLS2 S-acylation delays endocytosis and stabilises the FLS2-BAK1 complex at the plasma membrane, thereby helping to sustain signalling competence. This failure to sufficiently prolong signalling competence also explains the defects observed in *fls2c/proFLS2:FLS2 C^1132,1135^S* plants where subsequent signalling outputs such as *PR1* induction, growth inhibition and, ultimately, resistance to pathogenic bacteria (figure 3) are greatly impaired. Following activation, FLS2 is endocytosed and degraded, with new FLS2 being synthesised within approximately 1 hour of initial flg22 perception [38, 39]. Our observation that FLS2 S-acylation returns to near basal levels after 1 hour correlates with reported timings of degradation and *de-novo* FLS2 synthesis [38] but, at present, we cannot exclude an active process of FLS2 de-S-acylation prior to endocytosis.

Sequence analysis of RKs from across the Streptophyte lineages indicate that the S-acylation site identified here at the C-terminus of the FLS2 kinase domain is conserved across plant RK families throughout evolutionary history (figure S2). Assessment of the elongation factor-Tu perceiving receptor kinase EFR indicates that, similarly to FLS2, it undergoes ligand responsive S-acylation at this conserved cysteine (Cys 975). Mutation of this cysteine in EFR recapitulates the downstream signalling defects observed in S-acylation defective FLS2. We therefore hypothesise that there is a conserved role for S-acylation at these sites in other plant RKs. Recently, the P2K1/DORN1/LecRK-I.9 RK was proposed to undergo de-S-acylation followed by re-S-acylation during immune responses [40]. However, the site proposed is unique to the LecRK family, being distinct in proposed function, location, sequence, and structure to the universally conserved cysteine identified here that is also present in P2K1 but was not considered in the previous work. These data demonstrate that, in common with other post-translational modifications, S-acylation may affect multiple sites within an RK with differing effects on RK function (e.g. this work and [25]). The position and effect of the S-acylation site identified here at the C-terminus of the FLS2 and EFR kinase domains is highly conserved amongst plant RKs, and is also found in the closely related receptor-like cytoplasmic kinases (RLCKs) that act downstream of activated RKs. This opens up the exciting possibility that S-acylation at the conserved C-terminal kinase site may potentially regulate the function of all RKs (and RLCKs) across plants in a similar manner to FLS2 and EFR. However, this hypothesis awaits further empirical testing.

RK signalling is initiated by binding of a ligand (e.g., flg22) to its receptor (e.g., FLS2), which then facilitates the binding of a co-receptor (e.g., BAK1/SERK3). While this constitutes the minimal ligand recognition complex, substantial evidence supports a far larger number of proteins being intimately associated with both unstimulated and activated receptors and co-receptors. Indeed, existing data indicates that during the process of activation RKs recruit or eject specific proteins from their complexes [13, 18, 41, 42], but precise molecular mechanisms determining these changes are not known. Live cell imaging of unstimulated FLS2 and BAK1 indicates that presence or absence of the RK FERONIA (FER), has marked effects on nanoscale organisation and mobility of RKs in the plasma membrane. In addition, activation of the RK FERONIA (FER) by its ligand RALF23 alters BAK1 organisation and mobility [43]. This indicates that both complex composition, and the activation state of individual components, affects behaviour of the whole complex. Changes in direct protein-protein interaction can be explained by allosteric effects. However, it is also possible that alteration of the immediate (annular) lipid environment composition, curvature, or structure, brought about by changes in the physical properties of the complex, would act to recruit or exclude proteins based on their solubility and packing in the membrane environment surrounding the complex. This is, in essence, one of the principles proposed to underlie the formation of membrane nanodomains [44, 45]. Activation of FLS2 following flg22 perception has been reported to decrease overall plasma membrane fluidity and increase plasma membrane order [46], while changing sterol abundance in the plasma membrane affects all stages of FLS2 signalling [47]. This indicates that membrane composition and structure have profound effects on receptor complex function and supports the principle of protein-lipid interactions affecting or effecting RK function. S-acylation, being a fatty acid-based modification of proteins, has been shown to affect protein physical character and behaviour in membrane environments [33, 48]. S-acylation also affects membrane micro-curvature [23], a key theoretical determinant of membrane component partitioning required for nanodomain formation [44]. We therefore hypothesise that FLS2 S-acylation modulates interactions between FLS2 and immune complex components and/or FLS2 proximal membrane lipid components and may effect changes in the composition of both. Altogether our data supports a model where flg22-induced, BAK1-dependent FLS2 S-acylation sustains FLS2-BAK1 association, prevents premature internalisation of activated FLS2 complex and, overall, acts to promote immune signalling.

## Acknowledgments

We would like to thank Antje Heese and Paul Birch for critical discussions and advice during the preparation of this manuscript. Ari Sadanandom provided *P. syringae* pv. tomato DC3000. *bak1-4* seed was provided by Delphine Chinchilla, *chc2-1* seed by Antje Heese and *pub12/13* seed by Libo Shan. This work was supported by BBSRC EASTBIO-DTP studentship (grant number BB/M010996/1) to SM and PH, BBSRC grants BB/M024911/1 and BB/P007902/1 to PH, Royal Society Grant RG140531 to PH, a Heisenberg fellowship from the Deutsche Forschungsgemeinschaft to SR, the Gatsby Charitable Foundation, the University of Zürich, the European Research Council (Grant Agreement 773153 IMMUNO-PEPTALK) to CZ, the European Molecular Biology Organization (EMBO Long-Term Fellowship 438-2018), and the German Research Foundation (DFG grant CRC1101-A09) to JG. SJ was supported by the Scottish Government’s Rural and Environment Science and Analytical Services division (RESAS).

## Author Contributions

### CRediT statement

**CHH**: Conceptualization, Methodology, Validation, Formal analysis (Equal), Investigation (Lead), Data curation (Equal), Writing - Review & Editing, Visualization. **DT**: Methodology, Validation, Investigation (Equal), Writing - Review & Editing. **SM**: Validation, Investigation. **MK**: Investigation. **SJ**: Methodology, Software, Investigation. **KX**: Investigation, Formal analysis (Equal). **SR**: Resources, Writing - Review & Editing, Supervision, Funding acquisition (Equal). **CZ**: Resources, Writing - Review & Editing, Supervision, Funding acquisition (Equal). **JG**: Methodology, Formal analysis (Equal), Investigation (Equal), Data curation (Equal), Writing - Review & Editing, Supervision, Visualization, Funding acquisition (Equal). **PAH**: Conceptualization (Lead), Methodology (Lead), Validation, Formal analysis (Lead), Investigation, Data curation (Equal), Resources, Writing - Original Draft (Lead), Writing - Review & Editing (Lead), Visualization (Lead), Supervision (Lead), Project administration (Lead), Funding acquisition (Equal).

## Materials and Methods

### Cloning and constructs

All *FLS2* mutant variants used in this study are based on fully functional *FLS2_pro_:FLS2* construct able to complement *fls2* mutants [49] containing the described FLS2 promoter and open reading frame with stop codon [50]. All construct manipulations were performed on pENTR D-TOPO based vectors. Nucleotide changes were generated using Q5 site directed mutagenesis kit (NEB) according to the manufacturer’s guidelines. *FLS2_pro_:FLS2-3xMYC-EGFP* and *FLS2_pro_:FLS2 C^1132,1135^S-3xMYC-EGFP* were made by recombinatorial cloning in yeast using a 3xMYC-EGFP PCR fragment amplified from *FLS2_pro_:FLS2-3xMYC-EGFP* [39] recombined with pENTR D-TOPO *FLS2_pro_:FLS2* or pENTR D-TOPO *FLS2_pro_:FLS2 C^1132,1135^S*. Entry clones were recombined into pK7WG,0 [51] using Gateway technology (ThermoFisher) to generate expression constructs. Expression constructs were transformed into *Agrobacterium tumefaciens* strain GV3101 pMP90 [52] for transformation of either Arabidopsis or *Nicotiana benthamiana*.

### Plant lines and growth conditions

All Arabidopsis lines were in the Col-0 accession background. The *fls2* [50], *bak1-4* [53], pub12/13 [10] and *chc2-1* [17] mutants have all been described previously. Transgenic *fls2/FLS2_pro_:FLS2* are already described [49] and *fls2/FLS2_pro_:FLS2 C^1132,1135^S* mutant variant lines were generated by Agrobacterium-mediated floral dip transformation [54]. T_3_ homozygous plants were used for all experiments. Plant material for experiments was grown on 0.5x MS medium, 0.8% phytagar under 16:8 light:dark cycles at 20 °C in MLR-350 growth chambers (Panasonic). For transient expression *Nicotiana benthamiana* plants were grown in 16:8 light:dark cycles at 24 °C and used at 4-5 weeks old. *A. tumefaciens* mediated transient expression was performed as described [55] using an OD600 of 0.1 of each expression construct alongside the p19 silencing suppressor at an OD600 of 0.1. Tissue was harvested 48-60 hours post infiltration.

### Eliciting peptides

Flg22 peptide (QRLSTGSRINSAKDDAAGLQIA) was synthesised by Dundee Cell Products (Dundee, UK). Elf18 peptide (Ac-SKEKFERTKPHVNVGTIG) was synthesised by Peptide Protein Research Ltd. (Bishops Waltham, UK).

### Seedling growth inhibition

For each biological replicate four days post-germination, 10 seedlings of the named genotypes were transferred to 12-well plates (5 seedlings per well), ensuring the cotyledons were not submerged. Wells contained 2 mL of 0.5x MS liquid medium with or without 1 μM flg22. Seedlings were incubated for 10 days and the fresh weight of pooled seedlings in each genotype for each treatment measured and an average taken. Flg22-treated/untreated weights for each genotype were calculated and presented data is an average of these data over three biological repeats. Fully independent biological repeats were performed over a period of 6 months with each genotype only being present once in each repeat.

### MAPK activation

Essentially as for [56]; 6 Arabidopsis seedlings of each genotype 10 days post germination were treated with 100 nM flg22 for the indicated times in 2 mL 0.5x MS medium. The 6 seedlings from each genotype at each time point for each treatment were pooled before further analysis. Fully independent biological repeats were performed over a period of 2 years with each genotype only being present once in each repeat. To assess EFR induced MAPK activation in *N. benthamiana* leaves from 5-week-old plants were transiently transformed by agrobacterium infiltration (OD600 0.1 of each construct plus p19 at OD600 0.1). 60 hours after transformation, 1 μM elf18 peptide in water or water only was infiltrated into the leaf and samples harvested after 15 minutes. Samples were subsequently processed as described [56].

### Reactive oxygen species production

Protocol based on Mersmann et al. (2010). Essentially, 10 seedlings of each genotype were grown for 14 days in 100 μL of 0.5x MS medium with 0.5% sucrose, in 96-well plates (PerkinElmer). Conditions were maintained at 22 °C with 12:12 light:dark cycles. Growth medium was exchanged for water with 10 nM flg22 for 1 hour, before replacing with water for a further 1 hour. ROS burst was then induced by replacing with a solution containing 100 nM flg22, 400 nM luminol (Fluka), and 20 μg/mL peroxidase (Sigma). Luminescence in each well was measured every 2 minutes in a Varioskan Lux (Thermo Fisher) for 30 cycles (approx. 1 hour total).

### Gene expression analysis

Ten seedlings of each genotype 10 days post-germination were treated with 1 μM flg22 or water for the indicated times. The 10 seedlings from each genotype/treatment at each time point for each treatment were pooled before further analysis. RNA was extracted using RNAeasy Plant kit with on column DNAse digestion according to the manufacturer’s instructions (Qiagen). Two micrograms RNA was reverse transcribed using a High-Capacity cDNA Reverse Transcription kit (Applied Biosystems). All transcripts were amplified using validated gene-specific primers [49]. Expression levels were normalized against *PEX4* (At5g25760) [57]. Each sample was analyses in triplicate (technical repeats) for each primer pair within each biological repeat. Relative quantification (RQ) was achieved using the ΔΔ_CT_ (comparative cycle threshold) method [58]. Significant differences between samples were determined from a 95% confidence interval calculated using the t-distribution. Fully independent biological repeats were performed over a period of 2 years with each genotype only being present once in each repeat.

### Bacterial infection assays

Infection assays of Arabidopsis lines by *Pseudomonas syringae* pv. tomato DC3000 were performed using seedling flood inoculation assays as described [59].

### Western blotting

FLS2 was detected using rabbit polyclonal antisera raised against the C-terminus of FLS2 as previously described [12, 60]. Anti-p44/42 MAPK (Erk1/2) (Cell Signalling Technology #9102) was used to detect phosphorylated MAPK3/6 according to manufacturer’s recommendations at 1:2000 dilution. Total Arabidopsis MAPK6 or *N. benthamiana* WIPK was detected using anti-Arabidopsis MPK6 (Sigma A7104) at 1:2000. Rabbit polyclonal antibodies against BAK1 were as described [35] or obtained from Agrisera (AS12 1858) and used at 1:5000 dilution. BAK1 phospho-S612 was detected using polyclonal rabbit antisera as described [36]. Plasma membrane H+ ATPase (Agrisra AS13 2671), Calnexin 1/2 (Agrisera AS12 2365) and UDP-glucose pyrophosphorylase (Agrisera AS05 086) were all used at 1:2500. HRP (ECL) or fluorophore (Licor CLx) conjugated secondary antibodies were used to visualise antibody reacting proteins, and Clean-Blot HRP (Thermo Fisher) secondary antibody was used for immunoprecipitation experiments. ECL Western blots were developed using SuperSignal West pico and femto in a 3:1 ratio by volume and signal captured using a Syngene G:box storm imager and quantitative photon count data stored as Syngene SGD files. Signal intensity was quantified from SGD files using Syngene GeneTools software. Fluorescent western blots were imaged using a Licor CLx controlled by ImageStudio and quantified using Licor ImageStudio.

### S-acylation assays

S-acylation assays using acyl-biotin exchange (ABE) were performed exactly as described [60]. For flg22-dependent changes in FLS2 S-acylation, 7 seedlings 10 days post germination were transferred to each well of 12-well plates. Each well contained 2 mL 0.5 × MS liquid medium. Seedling were incubated for 24 hours on an orbital mixer (Luckham R100/TW Rotatest Shaker, 38 mm orbit at 75 RPM). Thereafter, 100 μL of 0.5 × MS media containing flg22 was added to give a final flg22 concentration of 10 μM. Seedlings were incubated with continued mixing for the indicated times before harvesting. Relative S-acylation is calculated using: (EX+ intensity^SAMPLE X^ / LC+ intensity^SAMPLE X^) / (EX+ intensity^REFERENCE SAMPLE^ / LC+ intensity ^REFERENCE SAMPLE^) [61]. Sample X refers to the sample of interest, reference sample is typically untreated control plants.

### Co-immunoprecipitation assays using IGEPAL CA-630

Seedlings grown on solid 0.5x MS for 30-35 days were transferred to wells of a 6-well plates and grown for 7 days in 0.5x MS 2 mM MES-KOH, pH 5.8. Thereafter, the seedlings were transferred in beakers containing 40 mL of 0.5x MS 2 mM MES-KOH, pH 5.8 and subsequently treated with sterile mQ water with or without flg22 (final concentration of 100 nM) and incubated for 10 minutes. The seedlings were then frozen in liquid nitrogen and proteins extracted in 50 mM Tris-HCl pH 7.5, 150 mM NaCl, 10% glycerol, 5 mM dithiothreitol, 1% protease inhibitor cocktail (Sigma Aldrich), 2 mM Na_2_MoO_4_, 2.5 mM NaF, 1.5 mM activated Na_3_VO_4_, 1 mM phenylmethanesulfonyl fluoride and 0.5% IGEPAL for 40 minutes at 4 °C. Lysates were clarified at 10,000 g for 20 minutes at 4 °C and the supernatants were filtered through miracloth. For immunoprecipitations, α-rabbit Trueblot agarose beads (eBioscience) coupled with α-FLS2 antibodies [11] were incubated with the crude extract for 3 hours at 4 °C. Subsequently, beads were washed 3 times (50 mM Tris-HCl pH 7.5, 150 mM NaCl, 1 mM phenylmethanesulfonyl fluoride, 0.1% IGEPAL) before adding Laemmli buffer and incubating for 10 minutes at 95 °C. Protein samples were separated in 10% bisacrylamide gels at 150 V for approximately 2 hours and transferred into activated PVDF membranes at 100 V for 90 minutes. Immunoblotting was performed with antibodies diluted in blocking solution (5% fat-free milk in TBS with 0.1% (v/v) Tween-20). Antibodies used in this study: α-BAK1 [35] (1:5000); α-FLS2 [11] (1:1000); α-BAK1 pS612 [36] (1:3000). Blots were developed with Pierce ECL/ ECL Femto Western Blotting Substrate (Thermo Scientific). The following secondary antibodies were used: anti-rabbit IgG-HRP Trueblot (Rockland, 18-8816-31, dilution 1:10000) for detection of FLS2-BAK1 co-immunoprecipitation or anti-rabbit IgG (whole molecule)– HRP (A0545, Sigma, dilution 1:10000) for all other western blots.

### Co-immunoprecipitation assays using Diisobutylene-maleic acid (DIBMA)

For each genotype, 2 × 10 seedlings 10 days post-germination were transferred to each well of 12-well plate containing 2 mL 0.5 × MS liquid medium and incubated for 24 hours on an orbital mixer (Luckham R100/TW Rotatest Shaker, 38 mm orbit at 75 RPM). Thereafter, 100 μL of 0.5 × MS media containing flg22 was added to give a final flg22 concentration of 10 μM. The seedlings were further incubated with continued mixing for 20 minutes prior to harvesting and blotting dry. Tissue was lysed in 500 μl of lysis buffer (50 mM Tris-HCl pH 7.2, 10% v/v glycerol, 150 mM NaCl, 1% w/v DIBMA (Anatrace BMA101), with protease inhibitors (1% v/v, Sigma P9599)) and incubated at room temperature for 1 hour with gentle end-over-end mixing. The lysate was centrifuged at 5,000 g for 1 minute and the supernatant filtered through 2 layers of miracloth and combined with an additional 500 μl of filtered lysis buffer (without DIMBA). The clarified lysate was further centrifuged at 16,000 g for 1 minute and the supernatant applied to Amicon 0.5 mL 100 kDa MWCO spin filtration columns and centrifuged at 14,000 g until the retentate was <50 μl. The retentate was diluted to 500 μl with IP buffer (50 mM Tris-HCl pH 7.2, 10% glycerol, 200 mM L-arginine, with protease inhibitor (0.5% v/v, Sigma P9599) and centrifuged at 14,000 g until the retentate was <50 μl. The spin column was inverted and eluted into a 1.5 mL microfuge tube by centrifugation at 100 g for 1 minute The eluate was diluted to 500 μl with IP buffer, of which 20 μl was retained as an input control. Magnetic protein A beads (20 μl per IP reaction) were coated with 5 μg αFLS2 antibody overnight at 4 °C. The resulting beads were washed for 5 minutes with IP buffer containing 0.5 M NaCl followed by 2 washes with IP buffer and resuspended in IP buffer to 100 μl per IP reaction. The resulting FLS2-coated magnetic protein A beads were added to the DIBMA solubilised protein solution and incubated for 3 hours at room temperature with end-over-end mixing. Thereafter, the beads were washed three times with IP buffer, resuspended in 30 μl 2x LDS sample buffer with 2-mercaptoethanol and incubated at 65 °C for 5 minutes with shaking at 1000 RPM. The samples were separated on a 7.5% SDS-PAGE gel prior to transfer to PVDF and western blotting.

### Detergent resistant membrane preparation

To evaluate flg22-dependent changes in FLS2 detergent resistant membrane occupancy, 7 seedlings 10 days post-germination were transferred to each well of a 12-well plate, of which each well contained 2 mL 0.5 × MS liquid medium. Seedlings were incubated for 24 hours on an orbital mixer (Luckham R100/TW Rotatest Shaker, 38 mm orbit at 75 RPM), after which 100 μL of 0.5 × MS media containing flg22 was added to give a final flg22 concentration of 10 μM. The seedlings were further incubated with continuous mixing as before for 20 minutes before harvesting and snap freezing in liquid nitrogen. All subsequent steps were performed at 4 °C or on ice. The seedlings were then lysed in 0.5 mL ice cold 1% (v/v) IGEPAL CA-630 in 25 mM Tris-HCl pH 7.4, 150 mM NaCl, 2 mM EDTA, and 0.1% (v/v) protease inhibitors (Sigma-Aldrich, P9599). Lysates were clarified at 500 g and filtered through 1 layer of miracloth. The filtrate was centrifuged at 16,000 g for 30 minutes and the supernatant retained as a detergent soluble fraction (DSM) and mixed 3:1 with 4x reducing (2-mercaptoethanol) LDS sample buffer. The detergent resistant pellet (DRM) was gently washed with 1 mL lysis buffer, centrifuged at 16,000 g for 5 minutes, and the supernatant discarded. The resulting pellet was resuspended in 27 μL of 3:1 lysis buffer: 4x reducing LDS sample buffer, after which 25 μL of the DRM and DSM were separated by 7.5% SDS-PAGE and probed using anti-FLS2 polyclonal antibody as described [60]. Presence of PM H+ ATPase (DRM enriched), Calnexin 1/2 (DSM enriched) [62] and UDP-glucose pyrophosphorylase (cytosol) [63] were used as markers for DRM purity.

### Variable Angle - Total Internal Reflection Fluorescence (VA-TIRF) microscopy

VA-TIRF microscopy was performed using an inverted Leica GSD equipped with a 160x objective (NA = 1.43, oil immersion), and an Andor iXon Ultra 897 EMCCD camera. Images were acquired by illuminating samples with a 488 nm solid state diode laser, a cube filter with an excitation filter 488/10 and an emission filter 535/50 for FLS2-GFP, and a 532 nm solid state diode laser, a cube filter with an excitation filter 532/10 and an emission filter 600/100 for mRFP-REM1.3. Optimum critical angle was determined as giving the best signal-to-noise.

### Single particle tracking analysis

*Nicotiana benthamiana* plants (14-21 days old) were infiltrated with *Agrobacterium tumefaciens* (strain GV3101) solution of OD_600_ = 0.5 and imaged 24 to 30 hours post infiltration. Image acquisition was done within 2 to 20 min after 1 μM flg22 or corresponding mock treatment. For single particle tracking experiments, image time series were recorded at 5 frames per second (0.2 s exposure time) by VA-TIRFM. Analyses were carried out as previously described [64], using the plugin TrackMate7 [65] in Fiji [66]. Single particles were segmented frame-by-frame by applying a Laplacian of Gaussian filter and estimated particle size of 0.3 μm. Individual single particle were localized with sub-pixel resolution using a built-in quadratic fitting scheme. Single particle trajectories were reconstructed using a simple linear assignment problem [67] with a maximal linking distance of 0.2 μm and without gap-closing. Only tracks with at least seven successive points (tracked for 1.4 s) were selected for further analysis. Diffusion coefficients of individual particles were extracted using SPTAnalysis [68] based on cosine filtered and maximum likelihood estimates analysis of particles displacement.

### Co-localization analyses

*Nicotiana benthamiana* plants (14-21 days old) were infiltrated with *Agrobacterium tumefaciens* (strain GV3101) solution of OD = 0.2 and imaged 48 hours post infiltration. Images were recorded by VA-TIRFM using 250 ms exposure time. As previously reported [31], we emphasised cluster formation in the presented images by using the ‘LoG3D’ plugin [69]. Quantitative co-localization analyses of the FLS2-GFP and mRFP-REM1.3 were carried out as previously described [31], with minor modification. Using FiJi, images were subjected to a background subtraction using the “Rolling ball” method (radius = 20 pixels) and smoothed. We selected regions of TIRF micrographs with homogeneous illumination for both FLS2-GFP and mRFP-REM1.3. The Pearson co-localization coefficients were assessed using the JACoP plugin of FIJI [70]. For comparison, we determined values of correlation, which could be observed by chance by calculating the Pearson coefficient after flipping one of the two images.

### Structural modelling of FLS2 kinase domain

The FLS2 intracellular domain (amino acids 831-1173) was submitted to the Phyre2 [71] server (http://www.sbg.bio.ic.ac.uk/phyre2/) in default settings. The solved BIR2 kinase domain structure (PDB 4L68, residues 272-600) [72] was identified as the best match and FLS2 residues 841-1171 were successfully modelled onto the BIR2 structure (confidence 100%, coverage 89%). Cys to Ser mutational effects were modelled using Missense3D [73] in default settings.

---

**Supplemental figure 1.**
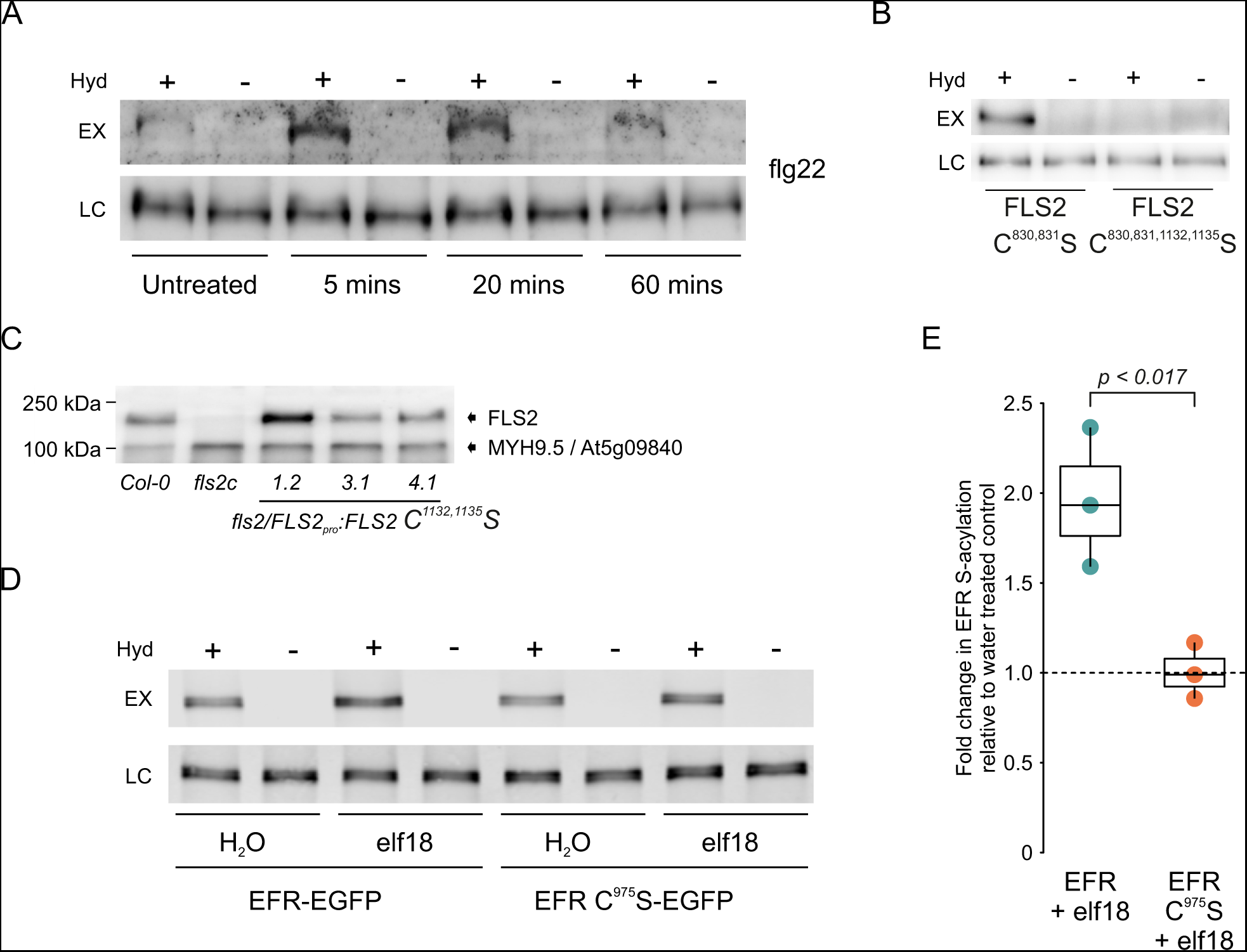
**A**. FLS2 C^830,831^S stably expressed in Arabidopsis *fls2* null mutant background retains the ability to be weakly S-acylated following flg22 treatment. S-acylation state was determined by acyl-biotin exchange assay. EX - indicates S-acylation state, LC - loading control, Hyd - indicates presence (+) or absence (−) of hydroxylamine. **B**. Mutation of FLS2 Cys1132,1135 to serine abolishes residual S-acylation observed in FLS2 C^830,831^S when over-expressed in *Nicotiana benthamiana*. EX - indicates S-acylation state, LC - loading control, Hyd - indicates presence (+) or absence (−) of hydroxylamine. **C**. Expression levels of FLS2 C^1132,1135^S in *fls2/FLS2_pro_:FLS2 FLS2 C^1132,1135^S* transgenic Arabidopsis lines used in this study. 50 mg total protein from 7-day old seedlings was loaded per lane. MYH9.5 / At5g09840 is a previously reported cross-reacting protein with the primary anti-FLS2 antibody used. **D**. EFR-GFP expressed in *Nicotiana benthamiana* undergoes S-acylation in a Cys975 dependant manner following 20 minutes of 1 μM elf18 treatment when. S-acylation state was determined by determined by acyl-biotin exchange assay. EX - indicates S-acylation state, LC - loading control, Hyd - indicates presence (+) or absence (−) of hydroxylamine. **E**. Quantification of EFR S-acylation state shown in D. elf18 induced changes to S-acylation state are shown relative to water treated (black dashed line). n = 3 biological repeats. Box plot shows median and IQR, whiskers indicate data points within 1.5 × IQR. Significance of difference between EFR and EFR C975S was determined by Student’s t-test.

**Supplemental figure 2.**
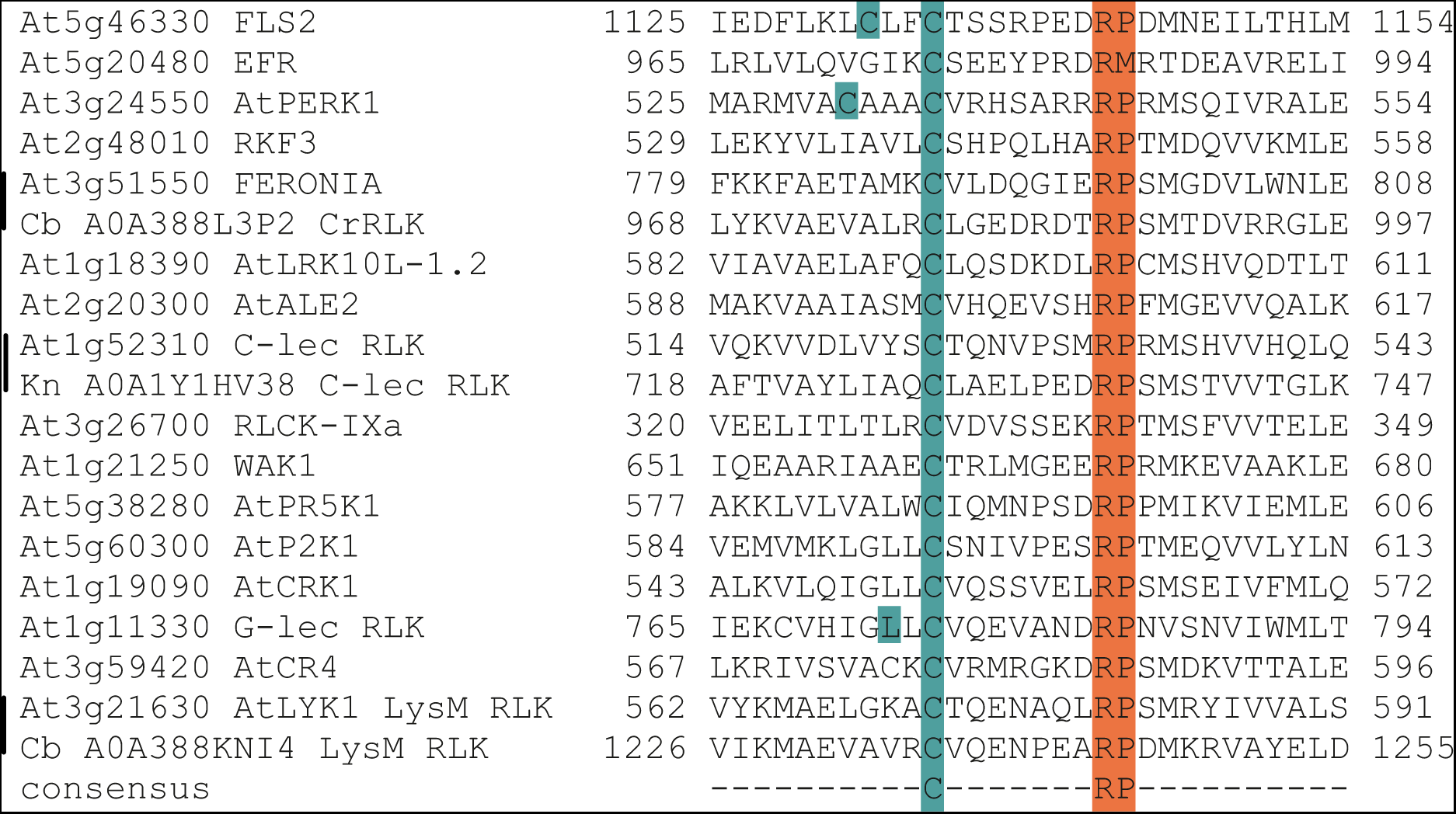
Receptor Kinases contain a conserved C-terminal cysteine within the kinase domain. Alignment using at least one representative member from each of the wider Arabidopsis RK superfamilies. Example receptor kinases found in *Chara braunii* (Cb) and *Klebsormidium nitens* (Kn) with clear sub-family members in Arabidopsis are also included as extant basal Streptophytes to illustrate evolutionary conservation of the proposed S-acylation site. Uniprot IDs are given for *Chara* and *Klebsormidium* sequences. Alignment is centred on the conserved C[X]_7_RP motif (orange) found in the loop between the G- and H-helices of the kinase domain. Putative S-acylation site cysteines are highlighted in teal with the conserved +7 RP motif in orange.

**Supplemental figure 3.**
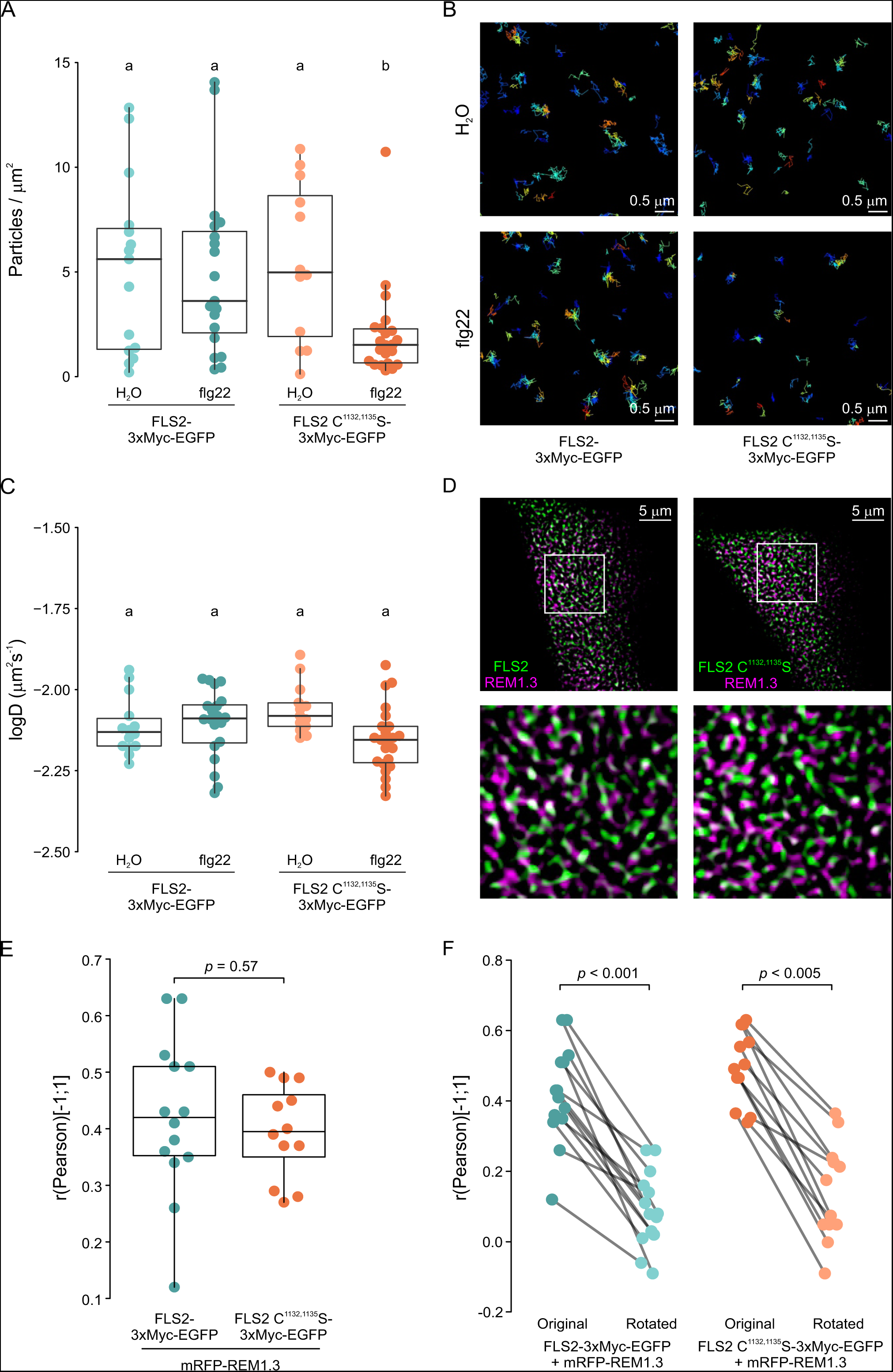
FLS2 S-acylation affects flg22 induced endocytosis but not unstimulated basal behaviour. **A**. FLS2-3xMyc-GFP and FLS2 C^1132,1135^S-3xMyc-GFP accumulate similarly when expressed in *N. benthamiana* in the absence of flg22, however, FLS2 C^1132,1135^S-3xMyc-GFP is cleared more rapidly than FLS2-3xMyc-GFP from the cell surface following flg22 exposure. Particle counts per μm2 at the plasma membrane of single cells using TIRF microscopy. Box plot shows median and IQR, whiskers indicate data points within 1.5 × IQR. FLS2-3xMyc-GFP mock n = 15 cells and 15076 particles, FLS2-3xMyc-GFP flg22 treatment n = 19 cells and 14717 particles, FLS2 C^1132,1135^S-3xMyc-GFP mock n = 12 cells and 12593 particles and FLS2 C^1132,1135^S-3xMyc-GFP flg22 treatment n= 22 cells and 7468 particles. p values calculated by ANOVA and confidence groups at p < 0.05 assigned using Tukey’s HSD test. **B**. Representative images from single particle tracking experiments of FLS2-3xMyc-GFP and FLS2 C^1132,1135^S-3xMyc-GFP at the plasma membrane using TIRF microscopy used to generate graph in A. Experiments were performed transiently in *N. benthamiana*. **C**. Quantification of average diffusion coefficient of single cells. Box plot shows median and IQR, whiskers indicate 1.5 × IQR. p values calculated by ANOVA and confidence groups at p < 0.05 assigned using Tukey’s HSD test. **D**. FLS2-3xMyc-GFP and FLS2 C^1132,1135^S-3xMyc-GFP form nanodomains in the plasma membrane and show similar co-localisation with mRFP-REM1.3 nanodomains when transiently expressed in *N. benthamiana* in the absence of flg22. Representative micrographs of FLS2-3xMyc-GFP and FLS2 C^1132,1135^S-3xMyc-GFP (green) co-localisation with mRFP-REM1.3 (magenta) at the plasma membrane of single epidermal cells using TIRF microscopy. **E**. Quantification of FLS2-3xMyc-GFP or FLS2 C^1132,1135^S-3xMyc-GFP co-localisation with mRFP-REM1.3 at the plasma membrane of single epidermal cells. FLS2-3xMyc-GFP n = 14 cells, FLS2 C^1132,1135^S-3xMyc-GFP n = 12 cells. Box plot shows median and IQR, whiskers indicate 1.5 × IQR. *p* value calculated using Student’s t-test. **F**. To determine whether measured co-localisation values shown in B (original) were significant, co-localisation analysis was repeated after rotation of the mRFP-REM1.3 image by 90 degrees (rotated). In all cases, co-localisation was reduced and overall significantly different, indicating that the co-localisation observed in B is both specific and significant. *p* values were calculated using Student’s t-test.

**Supplemental figure 4.**
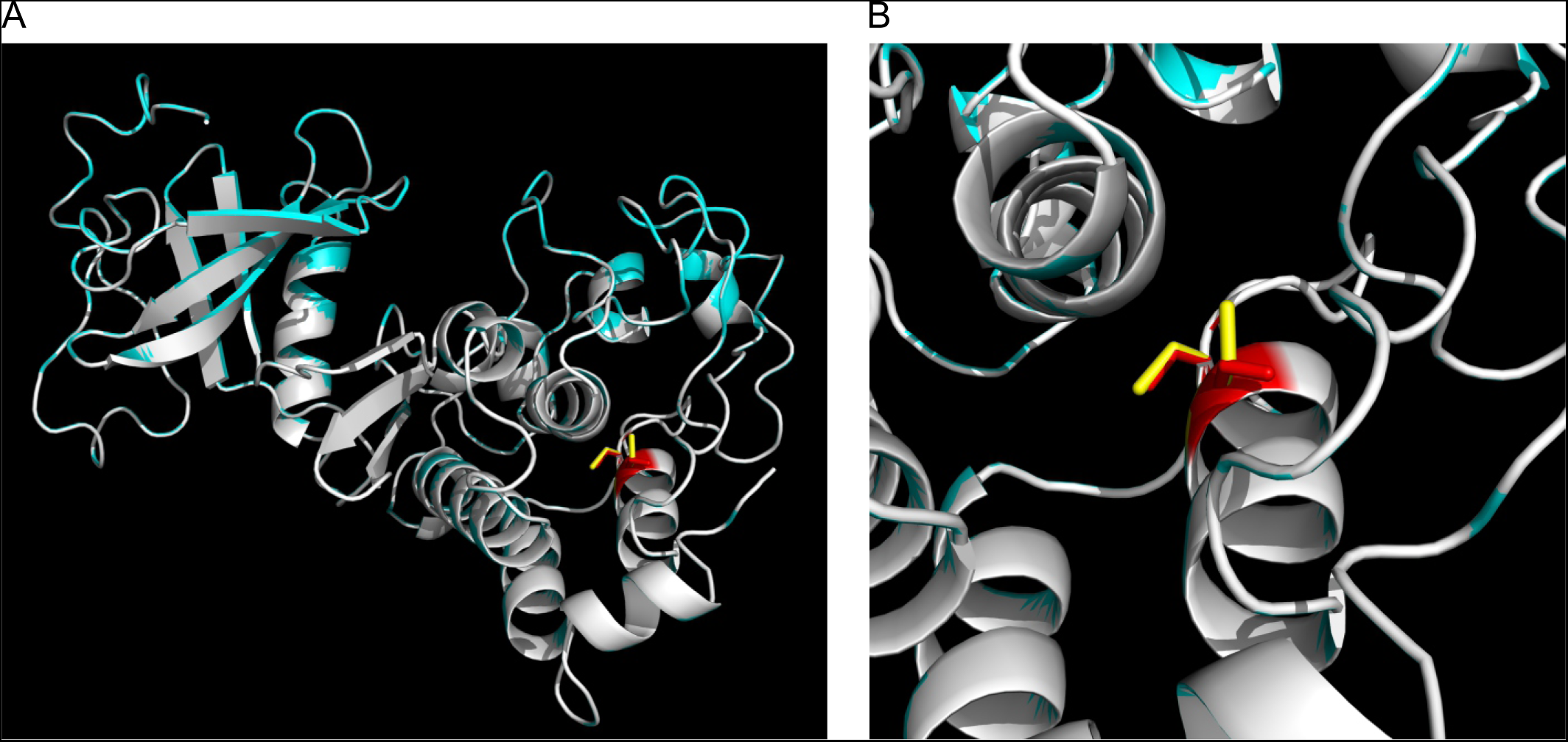
Mutation of kinase domain S-acylation site cysteines to serine in FLS2 is not predicted to affect kinase domain structure. **A**. Superimposition of the modelled structures of FLS2 (white) and FLS2 C^1132,1135^S (blue) kinase domains. **B**. Zoomed in view of Cys1132,1135 in FLS2 (yellow) and substituted serine (red) residues in FLS2 C^1132,1135^S. Only the proton of Ser1132 is predicted to diverge from the FLS2 structure, being rotated by ~110 degrees compared to the original cysteine. This rotation does not affect the position or packing of any other amino acid.

